# More from less: Genome skimming for nuclear markers for animal phylogenomics, a case study using decapod crustaceans

**DOI:** 10.1101/2020.12.05.413336

**Authors:** Mun Hua Tan, Han Ming Gan, Heather Bracken-Grissom, Tin-Yam Chan, Frederic Grandjean, Christopher M. Austin

**Author notes:** **Correspondence:** Mun Hua Tan.

## Abstract

Low coverage genome sequencing is rapid and cost-effective for recovering complete mitochondrial genomes for animal phylogenomics. The recovery of high copy number nuclear genes, including histone H3, 18S and 28S ribosomal RNAs, is also possible using this approach. In this study, we explore the potential of the genome skimming (GS) to recover additional nuclear genes from shallow sequencing projects. Using an *in silico* baited approach, we recover three additional core histone genes (H2A, H2B and H4) from our existing collection of low coverage decapod crustacean dataset (99 species, 69 genera, 38 families, 10 infraorders). Phylogenetic analyses based on various combinations of mitochondrial and nuclear genes for the entire decapod dataset and 40 species of crayfish (Infraorder Astacidea) found that the evolutionary rates for different classes of genes varied widely. The highlight being a very high level of congruence found between trees from the six nuclear genes and those derived from the mitogenome sequences for freshwater crayfish. These findings indicate that nuclear genes recovered from the same genome skimming datasets designed to obtain mitogenomes can be used to support more robust and comprehensive phylogenetic analyses. Further, a search for additional intron-less nuclear genes identified several high copy number genes across the decapod dataset and recovery of NaK, PEPCK and GAPDH gene fragments is possible at slightly elevated coverage, suggesting the potential and utility of GS in recovering even more nuclear genetic information for phylogenetic studies from these inexpensive and increasingly abundant datasets.

## Introduction

Outputs from second-generation sequencing (SGS) platforms enable the rapid and inexpensive assembly of animal and plant genomes (Austin et al., 2015, Austin et al., 2017, Tan et al., 2018a, Li et al., 2009, Colbourne et al., 2011). While this supports global collaborative efforts aiming to sequence an ever increasing diversity of species for tackling large-scale and ambitious biodiversity assessment and phylogenomic projects (Lewin et al., 2018, Consortium, 2013), for certain animal and plant groups with large and repetitive genomes, the problem of generating sufficient data for comprehensive phylogenetic and biodiversity-related studies can still be challenging.

Consequently, an increasing number of laboratories are using SGS to generate partial genome sequences at low cost (Gan et al., 2014) but with sufficient coverage to consistently recover highly-repetitive genes (Gan et al., 2018, Tan et al., 2018c) for potentially large numbers of individuals for phylogenetic analyses. These studies have mostly focussed on the recovery of genes from plastid genomes that are highly abundant in cells. While these genes are useful for taxonomic purposes, they can either be rapidly- or slowly-evolving and therefore limited in their power to consistently recover relationships over a range of evolutionary depths. Further, evolutionary relationships from mitochondrial genes have the limitation that they are linked and therefore represent just a single gene tree that may not reflect the true evolutionary history (Timm and Bracken-Grissom, 2015).

A relatively recent development that can help address these limitations is the discovery that repetitive nuclear genetic elements, predominately from the nuclear ribosomal cluster, can also be recovered from partial genome scans (i.e. genome skimming). The data derived by genome skimming (GS), defined as a method to obtain the high-copy fraction of a genome via shallow sequencing (Straub et al., 2012), contain numerous reads from repetitive nuclear genes (Besnard et al., 2016, Govindarajulu et al., 2015, Richter et al., 2015, Straub et al., 2012, Zimmer and Wen, 2015, Dodsworth, 2015, Malé et al., 2014). In recent studies on freshwater crayfish species, we showed that we can consistently recover genes from the nuclear ribosomal cluster (18S and 28S) and also a protein-coding gene (histone H3) (Grandjean et al., 2017, Tan et al., 2018c).

This study further explores the potential of GS using decapod crustaceans, a group with typically large genomes ranging from 1 to 40 Gbp, as proof of concept. Expanding upon Grandjean et al. (2017) and Tan et al. (2018c), we investigate the use of GS to recover non-traditional repetitive nuclear genes from a diverse range of decapod species (99 species, 69 genera, 38 families, 10 infraorders) and demonstrate the consistent recovery of three nuclear histone genes (H2A, H2B and H4), subsequently subjecting these to phylogenetic analyses focusing at the Infraorder level within the Decapoda and for a subset of samples representing freshwater crayfish (3 families, 14 genera and 40 species). Finally, we further evaluate the potential of GS by recovering several nuclear intron-less genes as well as finding known informative genes (NaK, PEPCK) (Chu et al., 2016) and the GAPDH housekeeping gene with slightly increased coverage.

## Material and Methods

A more detailed description of methods can be found in Supplementary Data 1.

### Genome skimming for four nuclear histones, 18S and 28S ribosomal RNA genes

This study used low coverage Illumina data obtained from previous projects (Tan et al., 2019, Tan et al., 2018b, Tan et al., 2018c, Tan et al., 2017, Tan et al., 2015) for 99 decapod species from 10 decapod infraorders within the suborder Pleocyemata, with the exclusion of the sister suborder, Dendrobranchiata (Supplementary Data 2). Genome skimming to extract histone (H2A, H2B, H3, H4) and ribosomal RNA (18S, 28S) genes was performed as previously described (Grandjean et al., 2017).

### Phylogenetic analyses and phylogenetic informativeness profiles

Maximum-likelihood phylogenetic analyses were conducted with MitoPhAST v3 (Tan et al., 2015) to construct trees based on datasets consisting of protein-coding genes (PCG) and ribosomal RNAs (rRNA) from mitochondrial (*mito*) and/or nuclear (*nuc*) origins: (I) 13 *mito* PCGs, (II) 2 *mito* rRNAs, (III) 4 *nuc* histones, (IV) 2 *nuc* rRNAs, (V) 13 *mito* PCGs + 2 *mito* rRNAs, (VI) 4 *nuc* histones + 2 *nuc* rRNAs, (VII) 13 *mito* PCGs + 2 *mito* rRNAs + 4 *nuc* histones + 2 *nuc* rRNAs. Analyses were nucleotide-based since all four histone genes show minimal variation at the protein level across all infraorders. Further, using a dataset of only protein-coding genes, we constructed an ultrametric tree (Bouckaert et al., 2014) and used PhyDesign (López-Giráldez and Townsend, 2011) to obtain profiles of phylogenetic informativeness for each gene.

### Preliminary scan for intron-less nuclear genes

A set of 640 intron-less nuclear genes was downloaded from the Intronless Gene Database (IGD) (Louhichi et al., 2011) as target gene candidates to simplify the search strategy, since these lack exon-intron structures. Using open reading frames predicted from assemblies (≥ 50 amino acid characters), single-copy orthologs of intron-less genes were identified with OrthoFinder v2.2.7 (Emms and Kelly, 2015).

### Genome skimming applied on datasets of variable sequence depths

Data from a crayfish species (*Cherax quadricarinatus*, Tan et al. (2020)), with an estimated genome size of 5 Gbp, was subsampled to generate multiple short read datasets at sequencing depths of 0.1×, 0.5×, 1× and 2×. Using OrthoFinder v2.2.7 (Emms and Kelly, 2015) as previously described, these datasets were scanned for: (1) intron-less genes, (2) the housekeeping gene GAPDH, and (3) NaK and PEPCK typically applied to multiple decapod phylogenetic analyses (Tsang et al., 2008, Tsang et al., 2014, Chu et al., 2016).

## Results

### Genome skimming recovers additional histone genes

Raw sequences used in this study are available on the Sequence Read Archive (SRA) under BioProject PRJNA485382. Genome skimming recovered three new histone genes (H2A, H2B, H4) and the same H3, 18S and 28S rRNA genes recovered by Grandjean et al. (2017) for all decapod species with minor exceptions (Supplementary Data 2, 3). For five species, only partial 28S rRNA genes were recovered, but with large gaps, and so these were excluded from further analyses and only partial histone H1 were obtained for 25 samples. The average sequence lengths for each recovered histone gene corresponds to expected full-length sizes. Per-gene coverage analysis showed an average of ∼100× for the mitogenome but were generally much higher for the nuclear genes (∼300×). Fig. 1A illustrates pairwise sequence identity for each mitochondrial and nuclear gene, showing an overall higher sequence divergence for mitochondrial genes.

**Fig. 1.**
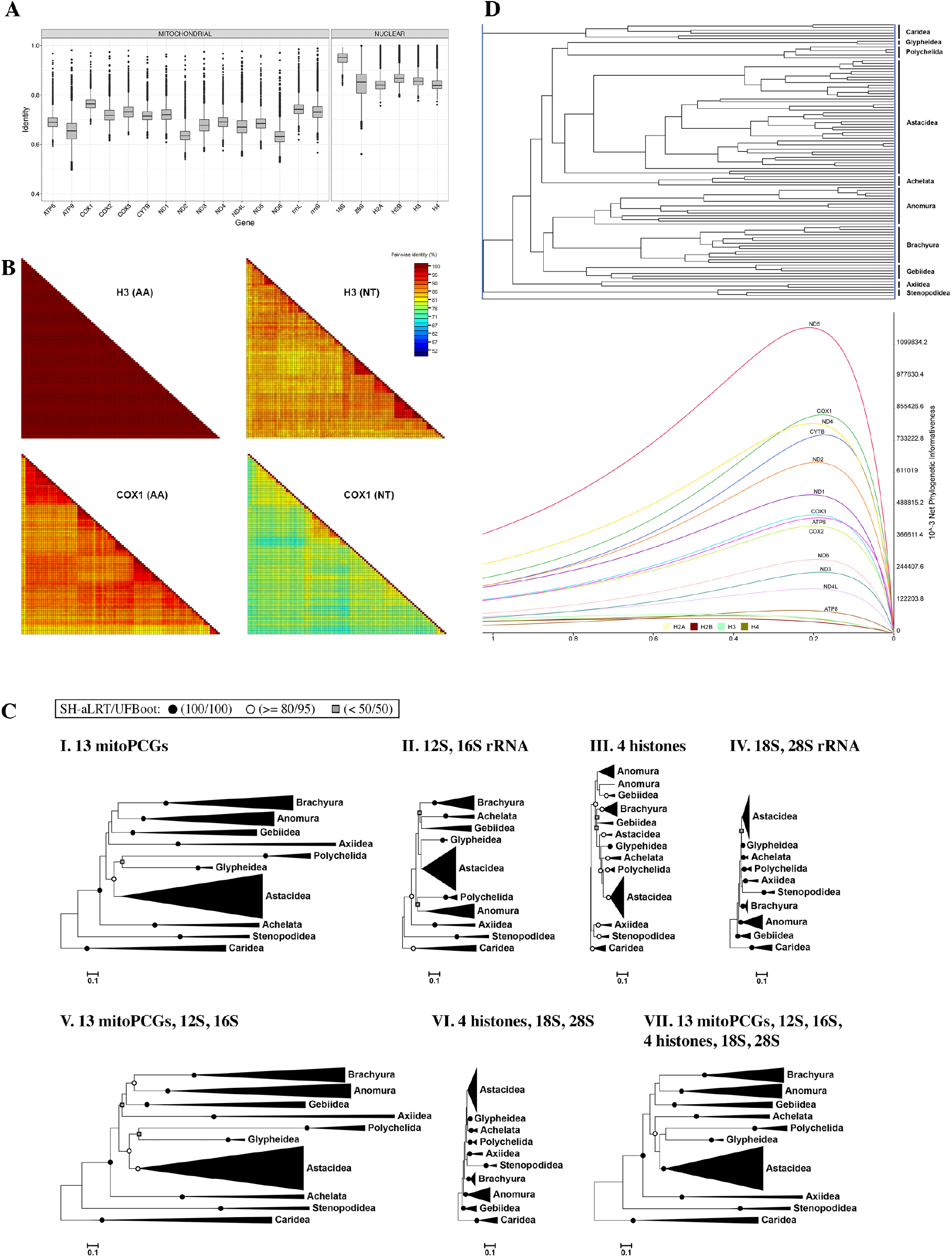
Sequence identities and phylogenetic analyses with mitochondrial and nuclear genes. **A**. Based on 99 decapod species, a higher overall divergence level is observed for mitochondrial genes than for nuclear genes. **B**. Matrices show pairwise sequence identities for the nuclear histone H3 compared to the mitochondrial COX1 at nucleotide and amino acid levels. **C**. Collapsed into decapod infraorders, branch lengths of every maximum-likelihood tree are adjusted to the same scale. High SH-aLRT / UFBoot support values are indicated with closed circles (100/100) and open circles (≥80/≥ 95). Unlabelled nodes have mid-level support values whereas weak nodes are labelled with grey rectangles (< 50 for either SH-aLRT or UFBoot). **D**. Phylogenetic informativeness profiles of 13 mitochondrial protein-coding genes and 4 nuclear histone genes.

Identity matrices in Fig. 1B and Supplementary Data 4 indicate overall highly conserved protein sequences for all four histones, with slightly higher variability observed for H2A and H2B genes. At the nucleotide level, histone sequences are relatively more divergent (average identities 83.0% to 85.4%) with noticeably greater levels of differentiation at higher taxonomic levels. In contrast, pairwise identities of COX1 at the amino acid level show obvious sequence differences (average identity 84.8%) with higher divergence levels when comparisons were done based on nucleotide characters (average identity 75.0%).

### Phylogenetic analyses

Maximum-likelihood phylogenetic trees from the seven datasets are shown in Fig. 2, with clades collapsed into infraorders. Rooted with the Caridea, tree topologies vary depending on the dataset. In all trees, there is generally good nodal support for each infra-ordinal clade represented by high nodal support values (SH-aLRT ≥ 80%, UFBoot ≥ 95%) (Fig. 1C). In contrast, nodes at the deeper level are mostly unsupported, especially in trees based on histones and/or nuclear rRNAs. Trees based on these nuclear-only datasets also have shorter deep internodes whereas trees based on mitochondrial datasets have relatively longer internodes and even longer branch lengths (Supplementary Data 5). In all phylogenetic informativeness (PI) analyses, the fast-evolving mitochondrial genes generally yielded the highest informativeness, whereas all four histone genes show the lowest utility for most of the tested time scale, slightly exceeding ATP8 in informativeness at deeper evolutionary divergence times (Fig. 1D, Supplementary Data 6).

**Fig. 2.**
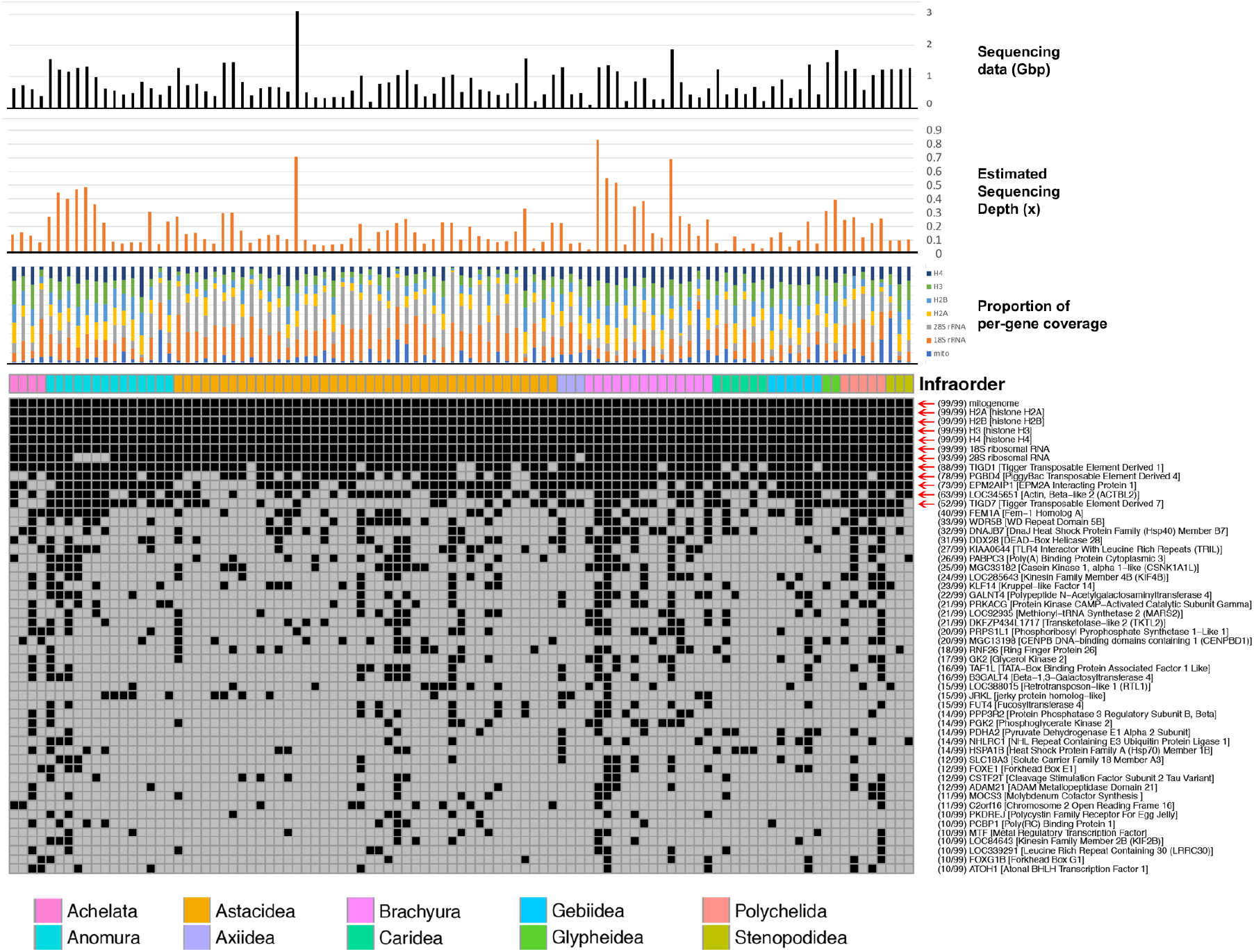
Presence of 4 nuclear histone genes, 18S rRNA, 28S rRNA and 45 other intronless protein-coding genes in 99 partial decapod genome scans. Black and grey squares represent the presence and absence of a gene, respectively. This figure reports single copy orthologs detected for 45 intron-less genes in at least 10 decapod datasets (in parentheses). Red arrows highlight genes detected in > 50% of datasets. Sequence depths are approximated based on genome sizes from the Animal Genome Size Database. For species lacking genome size information, genome sizes of different species from the next-closest genus, family or infraorder are averaged to provide the next best estimate of a genome size.

In contrast, relationships within the Astacidea infraorder, consisting of 40 crayfish species, show generally higher levels of congruence between the mitogenome and nuclear datasets (Supplementary Data 7). In general, the 28S and 18S data were more conserved, but did show some significant heterogeneity in evolutionary rates within and between lineages. The greatest congruence was between the combined nuclear dataset (bp = 6,629) and the mitochondrial datasets (shares 33/40 nodes with tree VII), with both superfamilies, all genera, and several major clades all recovered in the independent analyses. An analysis of the combined data for freshwater crayfish, represents one of the most comprehensive estimates of evolutionary relationships for this group based on taxon and gene sampling, comprising 12,120 bp of mitochondrial data and 6,629 bp of nuclear data (Tree VII, Supplementary Data 7).

### Scan for intron-less genes detects presence of high copy number genes

Fig. 2 displays the 45 intron-less genes detected, with those in majority of datasets indicated by long horizontal stretches of black squares and marked with red arrows. Most of these are genes often present in high copy numbers such as beta-actin-like protein (ACTBL2) or highly repetitive transposable elements including Piggybac (PGBD4) and various Tigger (TIGD1, TIGD7) subfamily genes. Histograms of sequence data (Gbp) and the estimated coverage (×) for each sample (from 0.02× to 0.83×) are also indicated.

### Increased sequence depths positively impacts the number of nuclear loci recovered

More intron-less genes were detected at elevated coverages in crayfish sample, with relatively better performance for the former (Table 1). For this dataset, partial NaK, PEPCK and GAPDH nuclear protein-coding genes were also detected in datasets with at least 1× coverage.

**Table 1.** **Recovery of nuclear gene fragments from the *Cherax quadricarinatus* crayfish (∼ 5 Gbp genome) dataset at variable sequencing depths**.

| Depth | Sequence data | Intron-less | NaK | PEPCK | GAPDH |
| --- | --- | --- | --- | --- | --- |
| 0.1× | 500 Mbp | 4 | - | - | - |
| 0.5× | 2.5 Gbp | 18 | - | - | - |
| 1.0× | 5 Gbp | 58 | ✓ | ✓ | ✓ |
| 2.0× | 10 Gbp | 87 | ✓ | ✓ | ✓ |
| **Max bp of gene recovered:** |  |  | 372 | 1458 | 885 |

## Discussion

This study uses decapod nucleotide datasets generated from low coverage Illumina sequencing of 99 species to investigate the potential of genome skimming (GS) for phylogenetic applications. The datasets were previously used for the recovery of mitogenomes, and in addition, we here demonstrate the consistent recovery of two nuclear rRNA genes and four histone genes across all major decapod groups. Three of the histone genes have scant representation in public databases (H2A, H2B, H4) and have not been previously reported from any GS study to our knowledge.

Many researchers are unaware of the treasure trove of potential genomic markers already available for their species or taxa of interest, be it in the form of public sequence databases (Leinonen et al., 2011) or as in-house generated datasets. While the higher throughput capacity of sequencing has enabled targeting genomic loci through sequencing-based approaches such as anchored hybrid enrichment (e.g. Wolfe et al. (2019) to yield large numbers of informative loci, these are still relatively expensive and require prior knowledge of genomic information to determine the targeted regions. On the other hand, the genome skimming approach can be applied to already-existing raw genomic sequence data previously generated for small-scale mitogenome or microsatellite marker projects. In addition, several protein-coding genes commonly used for phylogenetic studies can be recovered from higher-depth data. More importantly, the application of GS transcends the evolution of sequencing platforms, permitting one to recycle data generated not only from the more recent platforms (e.g. NovaSeq, HiSeq, MiSeq) but also older ones (e.g. Illumina GAIIx, SOLiD, Roche 454) (Besnard et al., 2016, Grandjean et al., 2017, Straub et al., 2012).

While GS seems promising for retrieving new genomic markers, it is necessary to identify the “sweet spot” at which sequence coverage can be minimised while accommodating for differences in genome size, the age or source of genetic material (e.g. tissue type, preservation method), systematic biases in sequencing (Tilak et al., 2018, Aird et al., 2011) and the relative abundance of mitochondria and nuclear copies among samples (Barazzoni et al., 2000, Herbst et al., 2017). In this study, total amount of data available for decapod species ranges from 93 Mbp to 3.4 Gbp, with an average of 807 Mbp. Independent of the data size, we observed variable levels of mapping coverage for the mitogenome and nuclear markers and while generally higher sequence depths were observed for nuclear histones and rRNA relative to the mitogenome, several datasets show contrary patterns

The subsampling of sequence data for the *Cherax quadricarinatus* crayfish dataset with known genome size has allowed us to minimally assess the effects of coverage on the recovery of different sets of genes (Table 1). Nuclear intron-less genes were easily detected in extremely low coverage datasets, this number substantially increasing with added sequence data, while NaK, PEPCK and GAPDH genes were consistently detected in datasets of at least 1× coverage. Based on this study, we recommend sequence coverage of about 2 to 3× as the optimal range for the GS approach, subject to adjustments according to the species and target genes of interest. The GS method could benefit from further improvement especially in the process of recovering genes, moving away from the use of seed sequences to other signature profiles and functional domains as identification, especially when targeting highly divergent genes.

It is frequently argued that mitochondrial genes are inappropriate when used as the sole marker for inferring evolutionary relationships (Hurst and Jiggins, 2005, Ballard and Whitlock, 2004, Bracken et al., 2009), with studies now including several nuclear markers (Bracken-Grissom et al., 2014, Bracken-Grissom et al., 2013, Grandjean et al., 2017, Schultz et al., 2009, Tsang et al., 2008, Tsang et al., 2014). Thus, our finding of a high degree of congruence between trees from mitochondrial sequences and nuclear sequences, especially from the expanded range of histone genes for the freshwater crayfish dataset is very noteworthy and bodes well for the greater use of genome skimming for phylogenetic and potentially molecular taxonomic studies. A caveat is that it will still be important to understand better the most appropriate taxonomic levels for maximum phylogenetic utility. This is illustrated by the finding that the addition of nuclear-based information at the infraorder level gave incongruent topologies and extremely short internodes, recalcitrant to any clear resolution of deeper relationships. This was further corroborated by the PI profiles of the four histone genes showing low utility across the whole tested time scale relative to that of mitochondrial protein-coding genes, regardless of codon positions considered. Our studies also indicate that the recovered nuclear genes, vary in their utility. The 18S and 28S gene sequences, commonly used in phylogenetics using traditional PCR methods, are the most conserved, but can show idiosyncratic levels of divergence in some species and lineages, leading to heterogeneity in branch lengths. Achieving consistent alignment is also problematics as nucleotide variation can vary widely along the molecular with insertions and deletion common. In contrast, the histone genes, being protein coding, are straightforward to align and show more even levels of divergence across the crayfish tree.

The infra-ordinal relationships within Decapoda remain unresolved and in need of ongoing investigation (Bracken et al., 2009, Shen et al., 2013, Tan et al., 2018b, Tan et al., 2015). While the histones, 18S and 28S genes recovered by GS in this study make available important additional markers for phylogenetic studies that can be combined with mitochondrial markers recovered from the same samples, they are insufficient for shedding new light on decapod relationships. However, our study indicates the potential for recovering additional protein-coding markers useful for studies of deep relationships among animal groups. With slightly elevated sequence coverage, GS can likely routinely recover additional protein-coding genes such as NaK and PEPCK, known to be informative in resolving decapod phylogenetic relationships (Chu et al., 2016, Tsang et al., 2008, Tsang et al., 2014).,

As the cost of sequencing continues to plummet, the prospect of GS as an effective tool for generating datasets for phylogenetics are increasing bright. Most of the samples used in this study were less than 1× coverage at a cost of several hundred dollars/sample. Using costing based on a NovaSeq (Deakin Genomics Centre) we estimate that over 5 Gb of data can now be generated for less than $200/sample and that with increased coverage, will consistently yield a greater range of nuclear markers that can be routinely used for phylogenetics.

## List of Supplementary Material

**Supplementary Data 1**. Detailed description of methods

**Supplementary Data 2**. Gene length and coverage for four nuclear histones, 18S and 28S ribosomal RNA genes recovered from 99 species from ten decapod infraorders

**Supplementary Data 3**. Sequence files for ribosomal RNA and histone genes

**Supplementary Data 4**. Pairwise identity matrices for COX1 and histone genes

**Supplementary Data 5**. Phylogenetic trees inferred from different combination of genes **Supplementary Data 6**. Phylogenetic informativeness profiles obtained for alignments with only codon positions 1+2 or 3

**Supplementary Data 7**. Phylogenetic trees with ultrafast bootstrap support values, inferred from different combinations of genes (expanded Astacidea clade)

### Acknowledgements

This work was supported by the Monash University Malaysia Tropical Medicine and Biology Platform and by the Deakin Genomics Centre.

