## Supplementary Data 1 for "More from less: Genome skimming for nuclear markers for animal phylogenomics, a case study using decapod crustaceans"

### Material and Methods

#### *Genome skimming for four nuclear histones, 18S and 28S ribosomal RNA genes*

This study used low coverage Illumina data obtained from previous projects (Tan et al., 2019) for 99 decapod species from 10 decapod infraorders within the suborder Pleocyemata (Supplementary Data 2), with the exclusion of the sister suborder, Dendrobranchiata. Genome skimming to extract histone (H2A, H2B, H3, H4) and ribosomal RNA (18S, 28S) genes was performed as previously described (Grandjean et al., 2017). Briefly, all four histone genes were recovered using contigs from either the *de novo* assembly or baited assemblies using sequences from closely-related organisms. Open-reading frames (ORF) were predicted from contigs with *getorf* (-minsize 30, -find 1 for nucleotide and -find 3 for amino acid) from EMBOSS v6.6.0 (Rice et al., 2000), followed by the identification of target genes based on sequence homology with *blastp* v2.6.0+ (Altschul et al., 1990).

#### *Nucleotide-based phylogenetic analysis workflow with MitoPhAST v3.0*

Since all four histone genes show minimal variation at the protein level across all infraorders (most sharing ≥ 90% identity), analyses were limited to only nucleotide-based datasets consisting of protein-coding genes (PCG) and ribosomal RNAs (rRNA) from mitochondrial (*mito*) and/or nuclear (*nuc*) origins:

I. 13 *mito* PCGs

II. 2 *mito* rRNAs

III. 4 *nuc* histones

IV. 2 *nuc* rRNAs

V. 13 *mito* PCGs + 2 *mito* rRNAs

VI. 4 *nuc* histones + 2 *nuc* rRNAs

***VII.*** 13 *mito* PCGs + 2 *mito* rRNAs + 4 *nuc* histones + 2 *nuc* rRNAs.

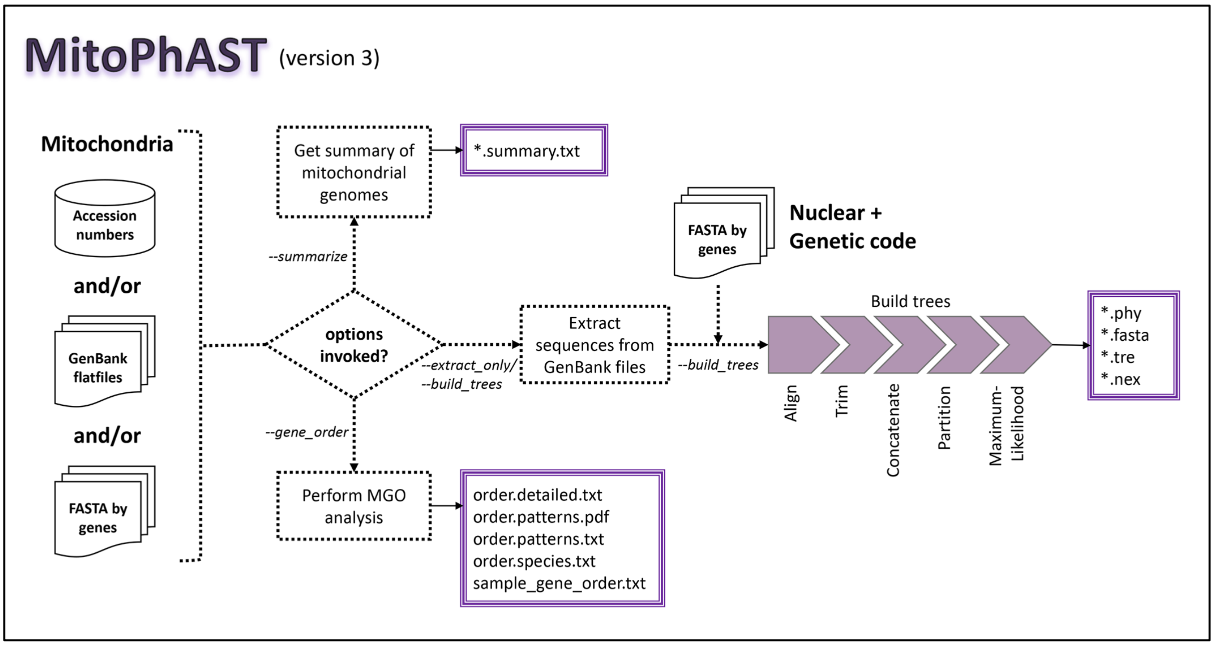


The version of MitoPhAST introduced here was used to construct Maximum-likelihood (ML) trees in this study (https://github.com/mht85/MitoPhAST, V3.0). While previous versions of MitoPhAST (Tan et al., 2018, Tan et al., 2015) perform only amino-acid based phylogenetic analyses, we have expanded the capabilities of the tool to include nucleotide-based ML analyses based on both mitochondrial and also nuclear genes. For each protein-coding gene, TranslatorX v1.1 (Abascal et al., 2010) is used to align nucleotide sequences based on their corresponding amino acid alignments. This is followed by trimming of ambiguously-aligned regions done internally with Gblocks v0.91 (Castresana, 2000), using less stringent parameters for typically shorter sequences. The MAFFT v7.394 aligner (Katoh and Standley, 2013) and Gblocks v0.91 trimmer (Castresana, 2000) is also used for non-coding genes sequences. In both cases, we advise that users manually check the alignments for potential alignment inaccuracies, or even to identify mis-annotations occasionally found in GenBank submissions. Multiple sequence alignments are then concatenated with FASconCAT-G v1.02 (Kück and Longo, 2014), generating super-alignments in both Fasta and Phylip formats. MitoPhAST then generates Nexus files specifying partitions by genes and codon positions and provides the files as input to IQ-TREE v1.5.5 (Nguyen et al., 2015), which further finds the best-fit partition schemes and models before constructing a Maximum-likelihood tree with SH-aLRT (Guindon et al., 2010) and ultrafast bootstrap (Minh et al., 2013) support values.

***BEAST analysis to obtain an ultrametric tree and phylogenetic informativeness profiles***

This was performed with mitochondrial and nuclear protein-coding genes. This was performed with mitochondrial and nuclear protein-coding genes. The jModelTest v.2.1.10 (Darriba et al., 2012) program was used to select appropriate nucleotide substitution models for each partition as site models in BEAST v.2.5.0 (Bouckaert et al., 2014). Since molecular evolution rates are variable across the different lineages, an uncorrelated, lognormal relaxed clock model was applied to each partition and the Yule Model was used as the tree prior since taxa in this study consist of individuals from different species. Caridean species were set as outgroup, since a large number of analyses generally place this infraorder at the base of Pleocyemata. Four BEAST MCMC runs of $5 \times 10^8$ were performed, sampling trees at every 5000 iteration. Convergence was visually checked for with Tracer v.1.7.1 (Rambaut et al., 2018) with 10% burn-in and checking for sufficient Effective Sample Size (ESS) (> 200). Logs and trees from the multiple MCMC runs were combined and a maximum clade credibility tree was constructed.

***Preliminary scan for intron-less nuclear genes***

A set of 640 intron-less nuclear genes was downloaded from the Intronless Gene Database (IGD) (Louhichi et al., 2011) as target gene candidates to simplify the search strategy, since these lack exon-intron structures. Using ORFs previously predicted from assemblies (≥ 50 amino acid characters), single-copy orthologs of intron-less genes were identified with OrthoFinder v2.2.7 (Emms and Kelly, 2015). Subsequently, the presence and absence of intron-less genes detected are visualized with the *pheatmap* R package (Kolde and Kolde, 2018).

***Genome skimming applied on datasets of variable sequence depths***

Data from a crayfish species (*Cherax quadricarinatus*, (Tan et al., 2020), with an estimated genome size of 5 Gbp, were subsampled to generate multiple short read datasets at sequencing depths of 0.1, 0.5, 1, 2 and 3×. Using OrthoFinder v2.2.7 (Emms and Kelly, 2015) as previously described, these datasets were scanned for: (1) intron-less genes, (2) the housekeeping gene GAPDH, and (3) NaK and PEPCK typically applied to multiple decapod phylogenetic analyses (Chu et al., 2016, Tsang et al., 2008, Tsang et al., 2014).
