## Supplementary figures and images for "More from less: Genome skimming for nuclear markers for animal phylogenomics, a case study using decapod crustaceans"

### Supplementary Data 4

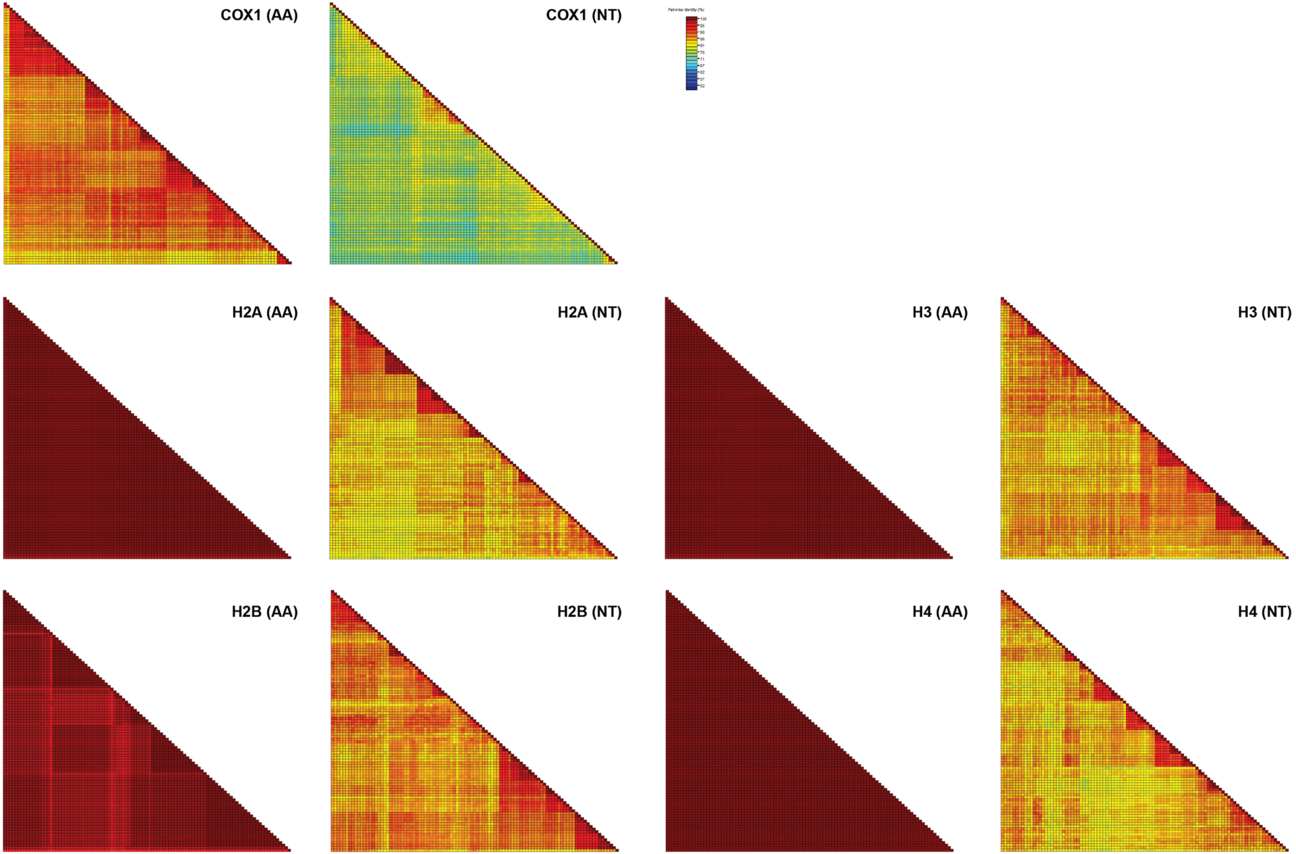

### Supplementary Data 6

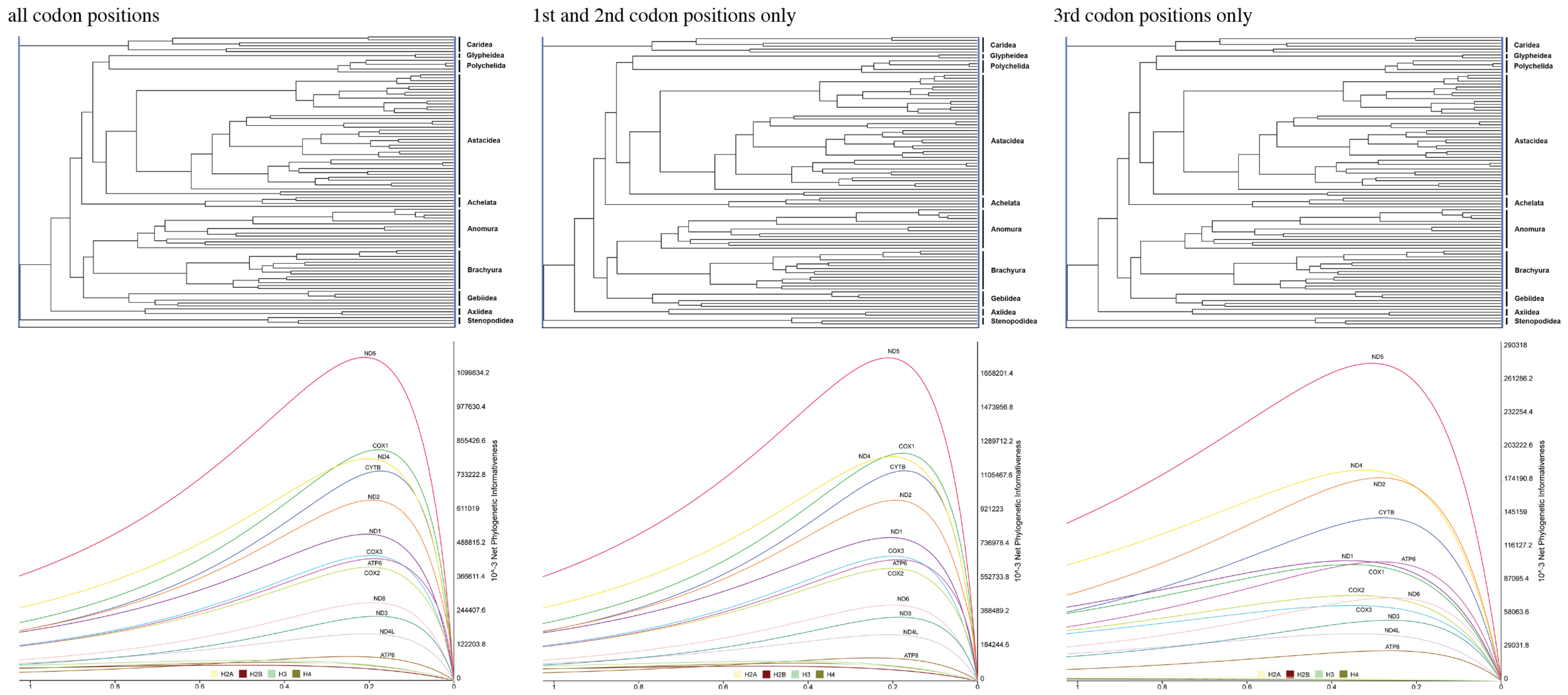
