## Supplementary Data 7 for "More from less: Genome skimming for nuclear markers for animal phylogenomics, a case study using decapod crustaceans"

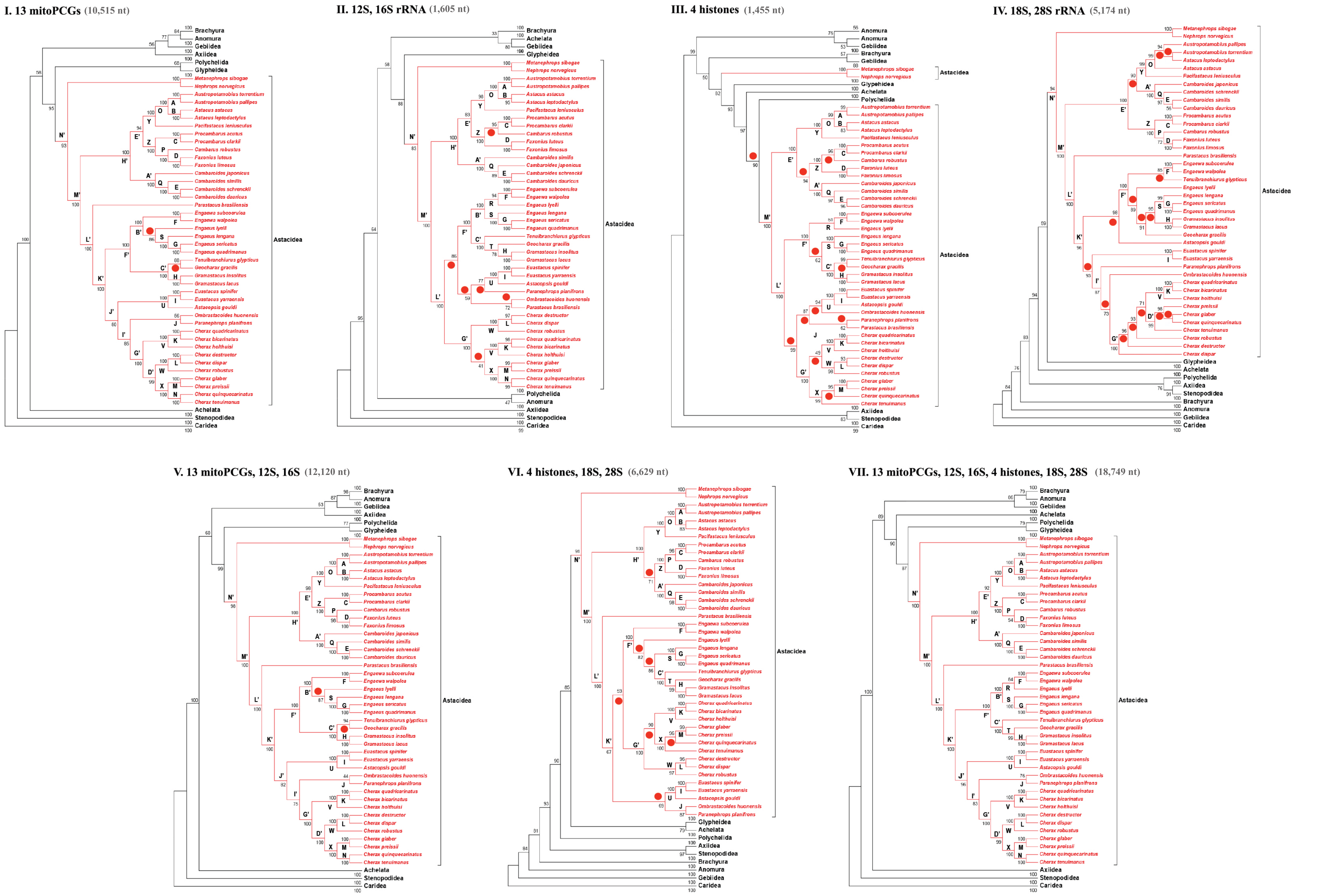
I. 13 mitoPCGs (10,515 nt)
II. 12S, 16S rRNA (1,605 nt)
III. 4 histones (1,455 nt)
IV. 18S, 28S rRNA (5,174 nt)
V. 13 mitoPCGs, 12S, 16S (12,120 nt)
VI. 4 histones, 18S, 28S (6,629 nt)
VII. 13 mitoPCGs, 12S, 16S, 4 histones, 18S, 28S (18,749 nt)
Astacidea
Brachyura
Anomura
Gebiidea
Axiidea
Polychelida
Glypheidea
Achelata
Stenopodidea
Caridea

Summary of commonalities in support for nodes within the Infraorder Astacidea and subordinate taxa for each tree represented in Figure 1C based on taxonomic classification and the tree based on the complete dataset. Infraorder refers to the Infraorder Astacidea; Superfamily refers to the superfamilies, Astacoidea and Parastacoidea; Family refers to the families Astacidae, Cambaridae[1] and Parastacidae; and Genus refers to the genera represented by more than 2 species, *Austropotamobius*, *Astacus*, *Procambarus*, *Faxonius*, *Cambaroides*, *Engaewa*, *Engaeus*, *Gramastacus*, *Euastacus* and *Cherax*. The "All Nodes" column represent the number of nodes within each tree common to the tree derived from the complete nucleotide data set (Dataset VII).

| Tree | Infraorder | Superfamily | Family | Genus | All Nodes |
|---|---|---|---|---|---|
| I. 13 mitoPCG | 1/1 | 2/2 | 2/3 | 10/10 | 38/40 |
| II. 12S,16SrRNA | 1/1 | 2/2 | 2/3 | 9/10 | 35/40 |
| III. 4 histones | 0/1 | 2/2 | 2/3 | 9/10 | 29/40 |
| IV. 18S,28SrRNA | 1/1 | 2/2 | 3/3 | 7/10 | 25/40 |
| V. 13mitoPCGs,12S,16 | 1/1 | 2/2 | 2/3 | 10/10 | 38/40 |
| VI. 4histones,18S,28S | 1/1 | 2/2 | 2/3 | 10/10 | 33/40 |
| VII. 13mitoPCGs,12S,16S,4histones,18S,28S | 1/1 | 2/2 | 2/3 | 9/10 | 40/40 |

[1] Follows the classification of Hobbs (1974) who includes the genus *Cambaroide*s within the Cambaridae, rather than Crandall and DeGrave (2017), who place representatives of the genus *Cambaroides* in the Family Cambaroididae.
